# Molecular insights on CALX-CBD12 inter-domain dynamics from MD simulations, RDCs and SAXS

**DOI:** 10.1101/2020.12.18.423531

**Authors:** Maximilia F. de Souza Degenhardt, Phelipe A. M. Vitale, Layara A. Abiko, Martin Zacharias, Michael Sattler, Cristiano L. P. Oliveira, Roberto K. Salinas

## Abstract

Na^+^/Ca^2+^ exchangers (NCX) are secondary active transporters that couple the translocation of Na^+^ with the transport of Ca^2+^ in the opposite direction. The exchanger is an essential Ca^2+^ extrusion mechanism in excitable cells. It consists of a transmembrane domain and a large intracellular loop that contains two Ca^2+^-binding domains, CBD1 and CBD2. The two CBDs are adjacent to each other and form a two-domain Ca^2+^-sensor called CBD12. Binding of intracellular Ca^2+^ to CBD12 activates the NCX but inhibits the Na^+^/Ca^2+^ exchanger of *Drosophila*, CALX. NMR spectroscopy and SAXS studies showed that CALX and NCX CBD12 constructs display significant inter-domain flexibility in the Apo state, but assume rigid inter-domain arrangements in the Ca^2+^-bound state. However, detailed structure information on CBD12 in the Apo state is missing. Structural characterization of proteins formed by two or more domains connected by flexible linkers is notoriously challenging and requires the combination of orthogonal information from multiple sources. As an attempt to characterize the conformational ensemble of CALX-CBD12 in the Apo state, we applied molecular dynamics (MD) simulations, NMR (^1^H-^15^N RDCs) and Small-Angle X-Ray Scattering (SAXS) data in a combined modelling strategy that generated atomistic information on the most representative conformations. This joint approach demonstrated that CALX-CBD12 preferentially samples closed conformations, while the wide-open inter-domain arrangement characteristic of the Ca^2+^-bound state is less frequently sampled. These results are consistent with the view that Ca^2+^ binding shifts the CBD12 conformational ensemble towards extended conformers, which could be a key step in the Na^+^/Ca^2+^ exchangers’ allosteric regulation mechanism. The present strategy, combining MD with NMR and SAXS, provides a powerful approach to select representative structures from ensembles of conformations, which could be applied to other flexible multi-domain systems.

**SIGNIFICANCE:** The conformational ensemble of CALX-CBD12, the main Ca^2+^-sensor of *Drosophila*’s Na^+^/Ca^2+^ exchanger, was characterized by a combination of MD simulations with SAXS and NMR data using the EOM approach. This analysis showed that this two-domain construct experiences opening-closing motions, providing molecular information about CALX-CBD12 in the Apo state. Ca^2+^-binding shifts the conformational ensemble towards extended conformers. These findings are consistent with a model according to which Ca^2+^ modulation of CBD12 plasticity is a key step in the Ca^2+^-regulation mechanism of the full-length exchanger.

## INTRODUCTION

High-resolution Nuclear Magnetic Resonance (NMR) spectroscopy is a well-established method to investigate structures of small proteins or protein domains in solution at atomic resolution. The major type of structural information obtained from NMR are short ^1^H-^1^H inter-nuclear distances (<6 Å) derived from the analysis of nuclear Overhauser effects (NOEs) in NOESY experiments (1, 2). Additional sources of structural information, such as residual dipolar couplings measured on weakly aligned samples (RDCs), rotational diffusion anisotropy determined from the analysis of ^15^N R_2_ and R_1_ relaxation rates, and paramagnetic relaxation enhancements (PRE) measured in samples labeled with a paramagnetic center, are particularly useful to the structural characterization of complexes and multi-domain proteins in solution (3–8). Small Angle X-ray Scattering (SAXS), on the other hand, is a low-resolution method and cannot provide information at atomic resolution (9). From the SAXS scattering profile it is possible to extract accurate structural parameters such as the average radius of gyration (R_*g*_), the degree of protein folding (Kratky plot) and the inter-domain orientation (“finger print”) from the pair distances distribution function, *p* (*r*), which is characteristic of the particle’s shape (10). Therefore, the information derived from SAXS is quite complementary to that obtained from NMR since they provide information in different length scales. Indeed, there are several examples of the simultaneous application of SAXS and NMR data to resolve the structure of large protein-protein complexes or multi-domain proteins (11–17).

Protein motions are essential for function (7, 18, 19). However, the structural characterization of flexible modular proteins imposes additional challenges. Under these conditions, a unique molecular envelope is not appropriate to describe SAXS data because different protein conformations may exist in solution (10, 20). The same is valid for NMR restraints such as RDCs, which are subjected to dynamic averaging due to motions up to the millisecond time scale (21, 22). In these cases, one possible approach is to build a large ensemble of different domain orientations representative of the protein dynamics and use the experimental data as a refinement filter of the ensemble (23, 24). In this strategy the principal concern is to avoid over-fitting, especially when an ensemble of structures is selected based on a single type of experimental data (23, 25, 26). The complementarity of NMR and SAXS data can be explored in order to break the degenerescence of the fitted parameters and prevent over-fitting (26, 27). Indeed, efforts have been made to calculate structural ensembles of multi-domain proteins when different inter-domain orientations co-exist in solution (5, 28), and many of them explore the synergy between NMR and SAXS data (16, 27, 29–32). One of the challenges in these approaches is to determine the right ensemble size, and to find the appropriate fractions of each model within the ensemble. A minimum representative ensemble may be obtained by optimization of the number of representative models and statistical weights using Bayesian inference (33, 34) or genetic algorithms (20, 35, 36). Alternatively, large ensembles may be calculated using clustering strategies followed by maximum-entropy refinement of the statistical weights of each cluster (37).

Na^+^/Ca^2+^ exchangers (NCX) are secondary active transporters that couple the import of Na^+^ to the extrusion of intracellular Ca^2+^. The NCX are essential for the maintenance of intracellular Ca^2+^ homeostasis, especially in excitable cells (38). They consist of a transmembrane domain, involved in ion translocation across the lipid bilayer, and a large cytosolic loop that is responsible for regulation of the exchanger activity by its substrates (39–41). The large intracellular loop contains a Ca^2+^-sensor, CBD12, which consists of two adjacent Ca^2+^-binding domains, CBD1 and CBD2, separated by a highly conserved linker of only three amino acids (Figure 1) (42, 43). The CBDs are, *β*-sandwich motifs, whose Ca^2+^ binding sites are located at the inter-strand distal loops (44–47). Therefore, the Ca^2+^-binding sites of CBD1 are located near the interface with CBD2 (Figure 1). CALX-CBD12 is the cytosolic Ca^2+^-sensor of the Na^+^/Ca^2+^ exchanger from *Drosophila*, CALX (48). While Ca^2+^ binding to CBD12 activates the mammalian exchangers, CALX is unusual because Ca^2+^-binding to CBD12 inhibits this exchanger (49). Studies of NMR spectroscopy of the CALX and the NCX-CBD12 constructs showed that the two CBDs are flexibly linked to each other in the Apo state, while binding of Ca^2+^ to CBD1 stabilizes a rigid and elongated inter-domain arrangement between CBD1 and CBD2 (50, 51). The flexibility of different NCX-CBD12 isoforms (NCX1-CBD12, NCX2-CBD12, NCX3-CBD12-AC and NCX3-CBD12-B) was characterized by SAXS using the ensemble optimization method (EOM) (20, 43, 52). It was found that a dynamic regime with a wide R_*g*_ distribution describes the Apo state SAXS profiles, while a narrower R_*g*_ distribution was selected in the presence of Ca^2+^. These observations are consistent with the greater inter-domain flexibility of the Apo state as indicated by NMR (50, 51). The same analysis has not been reported for CALX-CBD12.

**Figure 1:**
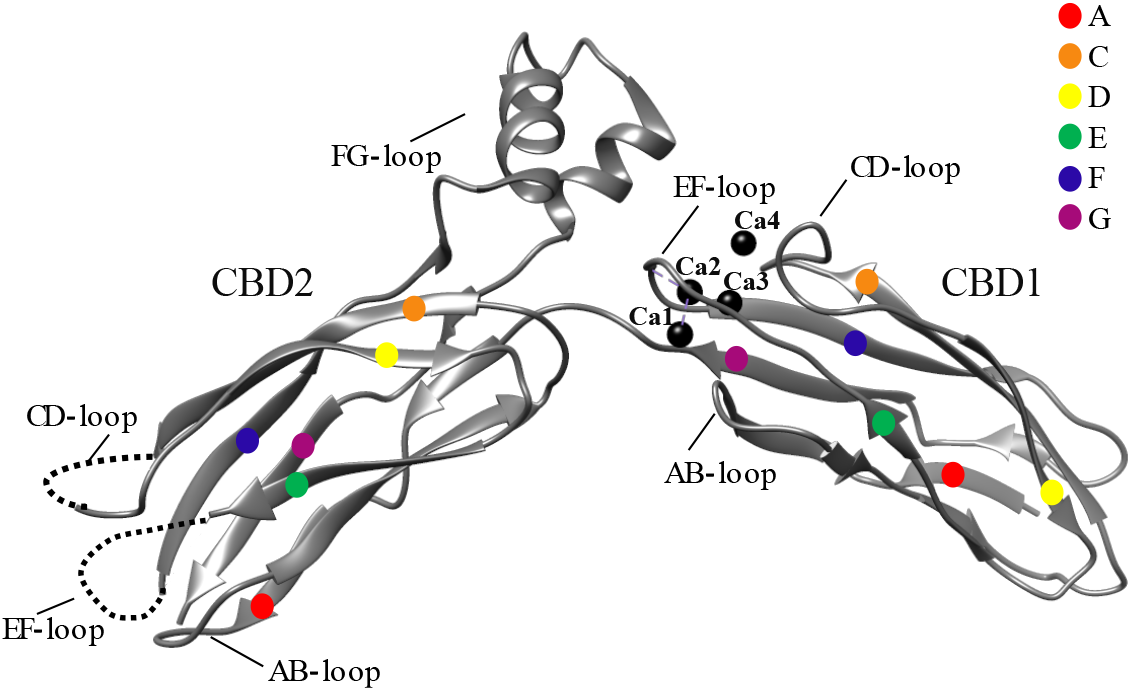
X-ray structure of the two-domain CALX-CBD12 construct (PDB 3RB5) in the Ca^2+^-bound state (42). Four Ca^2+^ ions at sites Ca1 – Ca4 in CBD1 are shown as black spheres. Segments with missing coordinates in the crystallographic structure are represented by segmented lines. The Ca^2+^-binding region of CBD1 corresponds to loops AB, CD and EF, and the inter-domain linker. The inter-domain interface consists of loops FG of CBD2 and EF of CBD1. The, *β*-strands are indicated according to the color legend.

Despite SAXS and NMR data indicate that CBD12 displays inter-domain flexibility (50, 51), an atomistic view of the dynamics of the Apo state, i.e. the various conformations and inter-domain arrangements that CBD12 adopts in solution, is still missing. Considering that atomistic description of these motions could give additional insights on the Ca^2+^ allosteric regulation mechanism of the CALX and the NCX exchangers, this question was addressed in this work using molecular dynamics (MD) simulations to generate an initial CALX-CBD12 ensemble, and the EOM approach (20) to select a minimal sub-ensemble that agrees with SAXS and RDCs data. In the absence of additional experimental data it is hard to tell that the selected conformations correspond to the native ensemble, however, they do offer atomistic insights on the kinds of motion experienced by CALX-CBD12 before Ca^2+^-binding. The strategy proposed here, combining MD simulations with SAXS and RDC data, might be useful for the analysis of other multi-domain systems that also experience significant inter-domain flexibility.

## MATERIALS AND METHODS

### CALX-CBD12 expression and purification

A DNA fragment corresponding to CALX1.1 residues 434–697 was PCR amplified from the CALX1.1 cDNA, and cloned in fusion with a N-terminal six-histidines tag in the pAE vector (53) using XhoI and EcoRI restriction sites. Protein expression was performed in *Escherichia coli* BL21-CodonPlus(DE3)-RIL (Agilent). Bacterial cell cultures were grown in LB at 37°C to an OD600 of 0.6, when the temperature was reduced to 18 °C and protein expression was induced by the addition of 0.4 mM of isopropyl-,*β*-D-thiogalactopyranosid (IPTG) for 18 hours. Cells were harvested by centrifugation, and cell pellets stored at −20°C. Approximately 10 g of cells were suspended in 200 ml of lysis buffer (20 mM Tris pH 7.0, 200 mM NaCl, 5 mM, *β*-mercaptoetanol, 0.1 µg/ml of pepstatin and aprotinin, 1.0 mM PMSF, and 1 mg/ml lysozyme), and subsequently lysed by sonication using a VCX 500 (Sonics) instrument. The cell lysate was clarified by centrifugation at 21000×g for one hour, the supernatant was applied into a 5 ml HisTrap affinity column (GE Healthcare) pre-equilibrated with buffer A (20mM sodium phosphate pH 7.4, 300 mM NaCl, 5 mM, *β*-mercaptoethanol, 10 mM Imidazole), washed with 50 ml of buffer A, and eluted during a gradient of 0.01 up to 0.5 M of Imidazole. Fractions containing CALX-CBD12 were combined, concentrated to 2 ml using an Amicon concentrator with a 3 kDa cutoff (Millipore), and applied into a Superdex 75 (16/600) gel filtration column (GE Healthcare) pre-equilibrated with the gel filtration buffer (20 mM Tris pH 8.0, 200 mM NaCl, 5 mM, *β*-mercaptoethanol, 1%(V/V) glycerol). Fractions containing CALX-CBD12 were combined and applied into a MonoQ 10/100 (GE Healthcare) ion-exchange column pre-equilibrated with the gel filtration buffer, and eluted in a gradient of 1.0 M of NaCl. Subsequently, fractions containing CALX-CBD12 were combined and buffer exchanged to 20 mM Tris pH 7.4, containing 5 mM of, *β*-mercaptoethanol, 200 mM of NaCl, 1% (V/V) of glycerol and 20 mM of EDTA. In order to prepare the SAXS samples, the EDTA was removed by buffer exchange using a 3 kDa Amicon centrifugation device. In the case of the Ca^2+^-bound sample, a suitable aliquot of a CaCl_2_ stock solution prepared in the same buffer as the protein was added just before the SAXS measurements. Protein concentration was determined by measuring the absorbance at 280 nm assuming *ϵ* _280_ = 25900 M^1^cm^1^.

### SAXS experiment

Small angle X-ray Scattering (SAXS) experiments were performed on a Xenocs-Xeuss laboratory SAXS equipment placed at the Institute of Physics of the University of São Paulo. This machine uses a microfocus source GENIX 3D with CuK*a* radiation, wavelength *λ*=1.5418 Å, Fox3D focusing mirrors and two sets of scatterless slits (Xenocs 2.0) for beam collimation. The scattering X-ray resulting from the sample-primary beam interaction was recorded on a 2D photon-counting Pilatus 300k detector at a sample-to-detector distance of 0.83 m. The scattering intensity was described as a function of the momentum transfer vector modulus *q* defined by *q*=4 *π* sin *θ*/*λ*, where 2 *θ* is the scattering angle. In the used setup the typical q range was 0.01<q<0.4 Å^−1^. SAXS data were collected on samples of CALX-CBD12 at a concentration of 4.2 mg.ml^−1^ (131 *μ* M) in the absence (Apo state) or in the presence of CaCl_2_ at a molar ratio of 6:1 (Ca^2+^: CBD12). Under these conditions, the four Ca^2+^-binding sites in the CBD1 domain must be fully occupied since the total protein concentration is significantly greater than the CALX-CBD12 Ca^2+^ dissociation constant measured by calorimetry, K_*d*_ = 1.6 M (42). CALX-CBD12 samples were injected in a Xenocs low noise flow cell and 900 s of exposures were recorded generating a total of 20 frames. Pairwise comparison of the collected frames indicated no radiation damage on the samples. The same arrangement was performed for an identically prepared sample but without the protein as a background and scattering-reference (plain water) sample to obtain the final intensity in an absolute scale. After the subtraction of the blank, all frames were averaged to obtain the final SAXS profiles. Fitting of the SAXS profile was carried out with GNOM (54) using the indirect Fourier transform (IFT) method to monodisperse systems. From this procedure the pair-distances distribution function *p* (*r*), the maximum diameter (D_*max*_), the radius of gyration (R_*g*_) and the intensity on *q* = 0 (*I* (0)) were determined. *I* (0) was used to calculate the protein molecular weight in solution (55). The calculation of the theoretical SAXS profile from known protein 3D coordinates was carried out using CRYSOL (56). A first assessment of CALX-CBD12 flexibility was performed applying the ensemble optimization method (EOM) (20, 36). RANCH (36) was used to build the initial CALX-CBD12 pool based on the crystal structure of the Ca^2+^-bound state (PDB 3RB5) (42) (Figure 1) and the protein amino acid sequence. Residues D550 – G555, in the linker between CALX-CBD1 and CALX-CBD2, were treated as flexible to generate a random pool of models, while the CBDs were treated as independent rigid bodies. A total of 10 000 models were built without symmetry or distance restraints. After the initial pool generation with RANCH, a total of 100 genetic algorithm (GA) cycles was performed with 1000 generations in order to select the optimal ensemble as implemented in GAJOE. As much as 50 ensembles were selected by GAJOE, each of them containing 20 or less models. Multiple copies of the same conformer in each ensemble were allowed during the optimization.

### Molecular Dynamics Simulation

Molecular dynamics trajectories (MD) of 3.4 *μ*s were calculated using the AMBER18 software package (57) at the target temperature of 300.0 K using the Berendsen weak coupling thermostat with 5 *p*s coupling time constant, pressure of 1.0 bar maintained with the Monte Carlo barostat, and the AMBER ff14SB force field (58). Starting structures for the simulations were taken from the crystallographic coordinates of CALX-CBD12 in the Ca^2+^-bound state (PDB ID 3RB5 chain A) (42). The four Ca^2+^ ions present at the CALX-CBD1 Ca^2+^-binding sites were removed in order to build the starting configuration for the Apo state. The missing coordinates in the crystallographic models were set up using the LEaP program utility from the AMBER-PDB preparation topology functions (59, 60). The two CALX-CBD12 initial structures (with and without the four Ca^2+^ ions) were solvated in explicit TIP3P water model using 10 Å dimension water layer around the protein in an octahedral box. To neutralize the box, Na^+^ and Cl^−^ ions were added as counter ions, additional Na^+^ and Cl^−^ ions were added to create a low salt concentration condition. First energy-minimization was performed during 2000 gradient steps, followed by heating up the system from 100 K up to 300 K within 50 *p*s in three stages with 25000 steps of 2 *f* s time step. All the analysis of structural parameters were carried out using CPPTRAJ routines (61). The inter-domain angle between CALX-CBD1 and CALX-CBD2 was measured considering three residues located near the centers of mass of CBD1 (A544), CBD12 (D551) and CBD2 (A688), and then the angle between the inter-residue vectors A554-D551 and D551-A688 was calculated for each MD trajectory snapshot. Structures were visualized with VMD (62).

### Residual Dipolar Couplings (RDCs) analysis

Experimental ^1^H-^15^N residual dipolar couplings (RDCs) measured on ^2^H/^15^N labeled CALX-CBD12 samples weakly aligned with Pf1 phages or mixtures of C12E5:n-hexanol (Apo state), and with Pf1 phages or compressed polyacrylamide gels (Ca^2+^-bound state), were previously reported (51). Fittings of the individual MD snapshots to the experimental ^1^H-^15^N RDCs were carried out using singular value decomposition (SVD) implemented in an in-house Python routine using NumPy (63, 64). Briefly, each snapshot was aligned applying CPPTRAJ routine (61) with respect to a reference frame taken at 400 ns using all non-hydrogen atoms, after stabilization of the simulation. The Cartesian coordinates of the N-H bond vectors were extracted, normalized, and used to calculate matrix **C**:

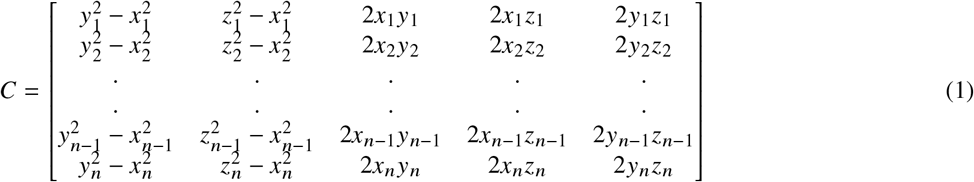

Order matrices, **A**, defined in an arbitrary Cartesian reference system were determined by solving a system of n linear equations (63, 64):

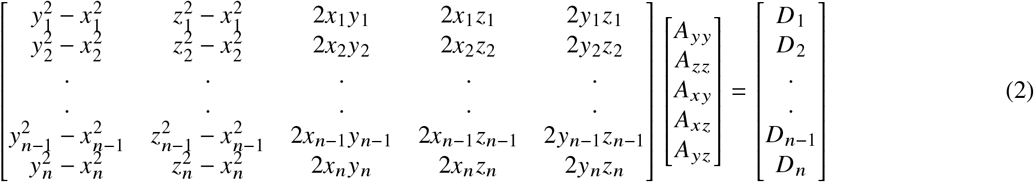

where *n* is the number of dipolar couplings,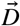 is the vector containing the experimental RDC values 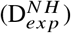, and A_*ij*_ are the five elements of the traceless order matrix, **A**. Once **A** was determined, RDC values 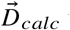 were back calculated according to 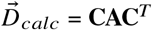 (63, 64). The agreement between 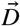 and 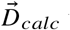 was evaluated using an R-factor (65). The smaller R-factor the better the correlation between 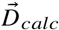 and 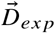, i.e. R = 0 means 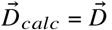. Alternatively, experimental ^1^H-^15^N RDCs were fitted to an ensemble of conformations as described by Niu and co-workers (66). In this analysis, population averaged N-H vector coordinates were calculated assuming *m* different conformations of CALX-CBD12 to build the new coordinates matrix **C** (Eq. 1) and an average alignment tensor for the ensemble. The six calculated terms ⟨*x*^2^⟩, ⟨*y*^2^⟩, ⟨*z*^2^⟩, ⟨*xy*⟩, ⟨*xz*⟩ and ⟨*yz*⟩ now correspond to population averages defined as

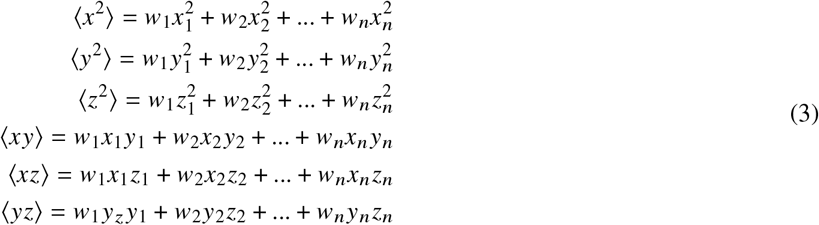

where *w*_*i*_ is the population weight of model *i* with 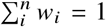. The fractions of the different models, *w*_*i*_, were allowed to fluctuate in order to minimize the R-factor using a Levenberg-Marquart algorithm implemented using in-house Python scripts and the Scipy library (67).

### Ensemble analysis combining MD with SAXS and NMR data

Two strategies were implemented to build a minimal structural ensemble of CALX-CBD12 using a Python routine outlined below (Fig. S1 in the Supporting Material). In the first approach, snapshots in the production phase of the MD trajectory (from 0.4 *μ* s up to 3.4 *μ* s) were saved every 0.2 ns, resulting in a pool of 15 000 models, which is considered a reasonable sampling size (36, 68) (Step 1A; 1st approach) (Fig. S1). This pool was used as an external library to SAXS-EOM (Step 2A; 1st approach) and the agreement of the EOM-selected representative conformers with experimental ^1^H-^15^N RDCs was evaluated by averaging the individual conformers coordinates (Step 3A, 1st approach) (Fig. S1). In the second approach, a larger pool of 54 000 models was built saving MD snapshots from the production phase of the trajectory every 60 *p*s (Step 1B; 2nd approach) (Fig. S1). The consistency of each snapshot with experimental ^1^H-^15^N RDCs was first evaluated by SVD, generating an R-factor distribution of the MD trajectory (Step 2B; 2nd approach) (Fig. S1). All models with R-factors lower than given thresholds were selected to form external pools to SAXS-EOM (Step 3B, 2nd approach) (Fig. S1). A total of six R-factor limits were tested for the Apo state (Pf1 data set): 0.54 (15 000 models), 0.53 (10 000 models), 0.52 (5 000 models), 0.47 (1 000 models), 0.43 (100 models) and 0.42 (50 models). Finally, the SAXS-EOM analysis (Step 4B, 2nd approach, Fig. S1) was carried out to select a minimum structural ensemble. CRYSOL (56) was used to compute the theoretical SAXS intensity of each individual model in the pool, and GAJOE, using the GA algorithm, to select the optimal ensemble that best describes the SAXS data (36). Briefly, a total of 100 genetic algorithm cycles (GA) was performed with 1000 generations in order to select the optimal ensemble, as implemented in GAJOE. As much as 50 ensembles were selected by GAJOE, each of them containing 20 or less models. Multiple copies of the same conformer in each ensemble were allowed during the ensemble optimization.

## RESULTS AND DISCUSSION

### CALX-CBD12 displays conformational heterogeneity in the absence of Ca^2+^

To investigate whether CALX-CBD12 SAXS scattering profiles are sensitive to the different inter-domain dynamics exhibited by the Apo and the Ca^2+^-bound states we recorded SAXS intensities of CALX-CBD12 samples prepared in the absence and in the presence of CaCl_2_ at a molar ratio of 1:6 (CBD12:CaCl_2_) (Fig. 2A). The pair-distance distribution function, *p* (*r*), observed for the Apo state, is characteristic of a slightly elongated molecule. In contrast, the *p r* obtained for the Ca^2+^-bound state shows an additional shoulder at ∼55 Å, which is characteristic of a two-domain protein with larger _*max*_ relative to the Apo state (Fig. 2B). Further analysis with the Kratky plot suggested that CALX-CBD12 behaves as a rigid body in the Ca^2+^-bound state, while the behavior in the absence of Ca^2+^ is consistent with two domains connected by a flexible linker (Fig. 2C) (69). The radius of gyration (*R*_*g*_) obtained from IFT analysis is 26.1 ± 0.1 Å and 28.3 ± 0.1 Å for the Apo and the Ca^2+^-bound states, respectively (Table 1).These results indicate the predominance of slightly less open structures in the Apo state, while extended conformations are predominant in the Ca^2+^-bound state. Since a crystal structure of CALX-CBD12 in the Ca^2+^-bound state is available (PDB entry 3RB5) (42) (Fig. 1), we compared the theoretical SAXS intensities with the experimental data using CRYSOL (56). The calculated SAXS intensities showed good agreement with the experimental data obtained in the presence of Ca^2+^ (*χ*^2^ = 2.3) (Fig. 2D), while a poorer agreement was observed in its absence, particularly at low *q* values and at the intermediate q range (*χ*^2^ = 10.2). Altogether, these observations are consistent with the greater flexibility of the Apo state relative to the Ca^2+^-bound state.

**Table 1:** CALX-CBD12 structural parameters obtained from the analysis of SAXS data recorded in the absence (Apo) and in the presence of Ca^2+^ (Ca^2+^-bound) at 6:1 molar ratio (CBD12: CaCl_2_).

| Sample | Radius of gyration - $R_g$ ($\text{\AA}$) | Maximum diameter - $D_{max}$ ($\text{\AA}$) | Molecular weight - MW (kDa) |
| --- | --- | --- | --- |
| Apo | $26.1 \pm 0.1$ | 87 | $33 \pm 3$ |
| $\text{Ca}^{2+}$ | $28.3 \pm 0.1$ | 93 | $33 \pm 3$ |
| 3RB5<sup>a</sup> | 28 | 93 | 30 |

<sup>a</sup> calculated values using CRY SOL (56) and the CALX-CBD12 crystal structure (PDB 3RB5 chain A).

**Figure 2:**
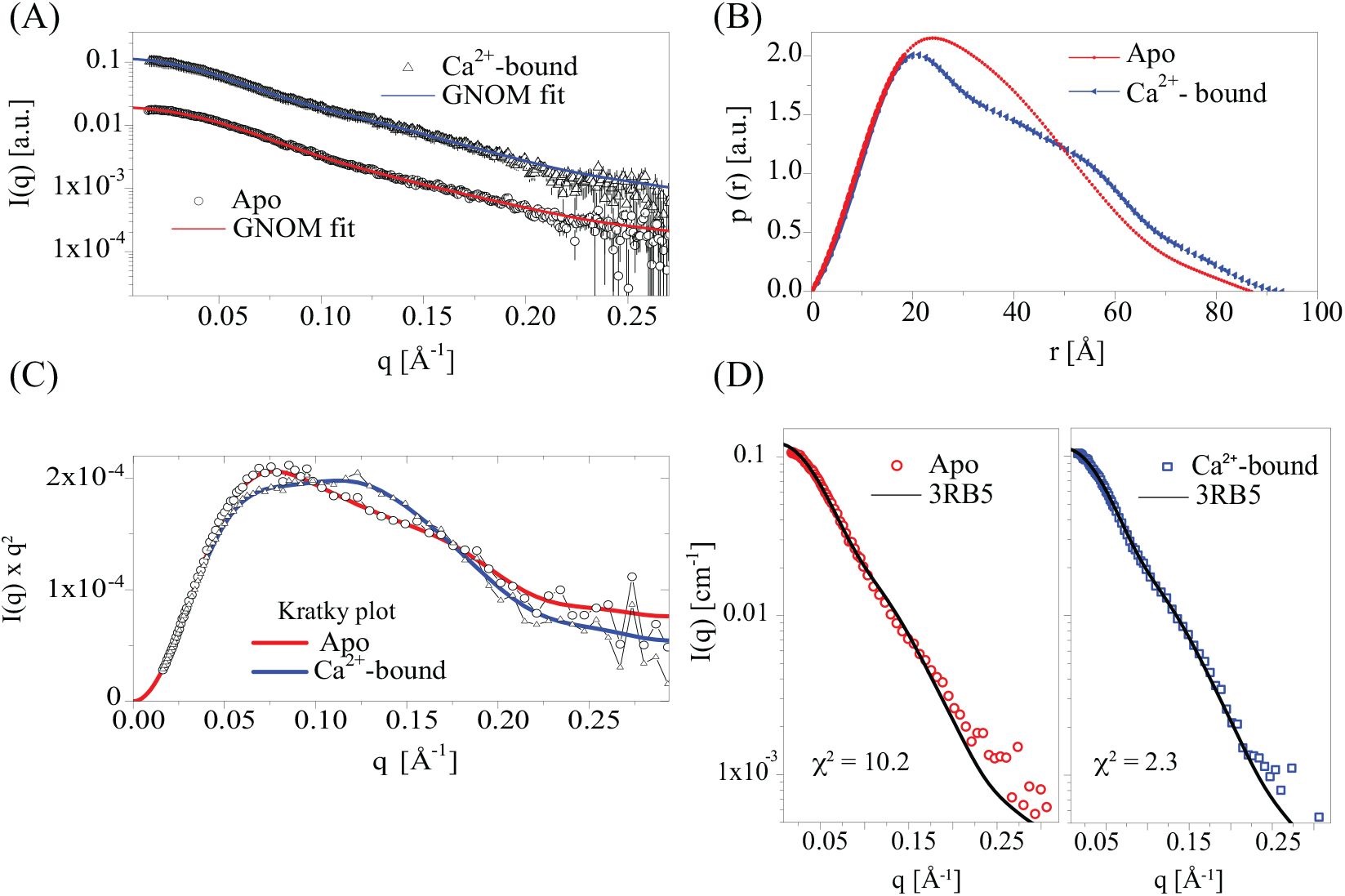
SAXS data analysis. **(A)** Experimental SAXS profiles of CALX-CBD12 in the absence (Apo) and presence of Ca^2+^ (Ca^2+^-bound) and IFT fits to the experimental data. (**B**) Pair distance distribution function, *p* (*r*), obtained for the Apo (red) and Ca^2+^-bound (blue) samples. (**C**) Kratky plot (*I* (*q*) ×*q*^2^ vs *q*) of CALX-CBD12 in the absence (Apo, red) and presence of Ca^2+^ (Ca^2+^-bound, blue). (**D**) Theoretical SAXS profiles calculated using the X-ray model (PDB 3RB5) (42) coordinates (solid line) overlaid on the experimental SAXS profiles of the Apo and Ca^2+^-bound states.

Analysis of the SAXS intensity profiles by rigid body modeling using the EOM approach (20) showed that two *R*_*g*_ distributions consisting of relatively compact models describe the Apo state SAXS intensities, while a single narrower *R*_*g*_ distribution with the CBDs in extended arrangements was selected for the Ca^2+^-bound state (**Fig. S2 in the Supplementary Material**). However, the agreement between calculated and experimental SAXS intensities in the Apo state remained poor (*χ*^2^ = 10.3), suggesting that the experimental data was not fully described by the EOM-selected ensemble (**Fig. S2-B**). This observation may be rationalized considering that the initial pool of CALX-CBD12 conformers was built using RANCH by randomization of the short inter-domain linker, while CALX-CBD1 and CALX-CBD2 were treated as rigid bodies. Hence, this initial pool may not capture the complexity of the conformational energy landscape of the two-domain construct. In order to start from a more realistic library of conformers, we carried out atomistic MD simulations of CALX-CBD12 in the Apo and in the Ca^2+^-bound states and used the MD ensemble as input for EOM.

### Sampling CALX-CBD12 energy landscape by atomistic MD simulations

Long 3.4 *μ*s MD trajectories of CALX-CBD12 in the Ca^2+^-bound and in the Apo states were calculated in explicit water starting from the CALX-CBD12 crystal structure as described in the Methods section. Analysis of the backbone RMSD relative to the first frame indicated that the simulations took approximately 0.4 *μ* s to equilibrate (Fig. 3A and 3B). The Ca^2+^-bound state deviated from the starting X-ray structure and was confined into a major conformational state characterized by RMSD of ∼ 5 Å and inter-domain angle ⟨*θ*⟩ = 81 ±7° during most of the trajectory, experiencing frequent jumps to other conformations with RMSDs in the range of 3.5 – 7 Å (Figs. 3A,B and **S3 in the Supplementary Material**). In contrast, the Apo state visited three major conformational states characterized by averaged RMSDs of approximately 3.5, 5.0 and 8.5 Å from the initial structure, and mean inter-domain angles ⟨*θ*⟩ = 116 ± 8, 87 ± 6 and 81 ± 5° (Figs. 3A,B and **S3**).

**Figure 3:**
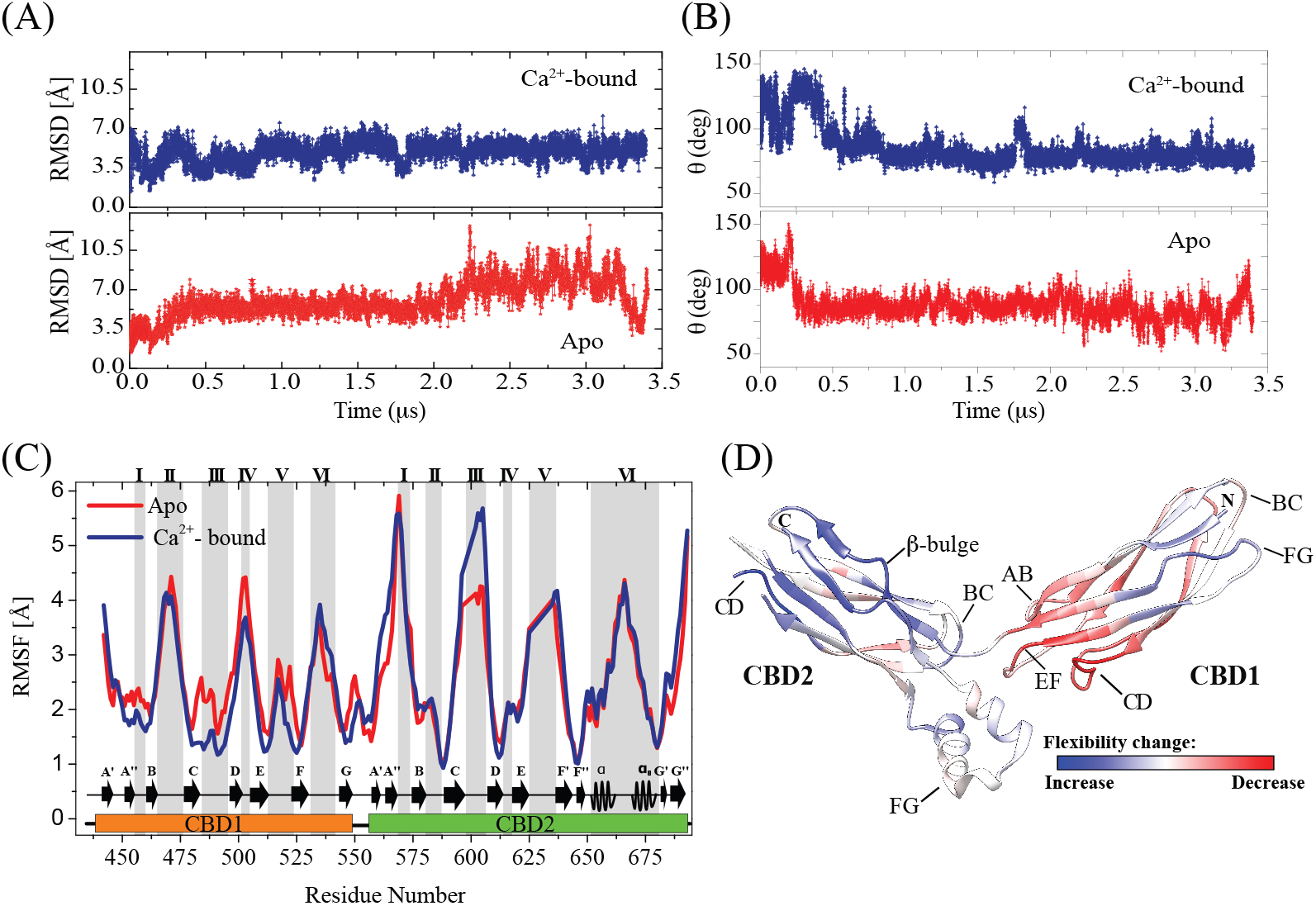
Analysis of molecular dynamics trajectories of CAXL-CBD12 in the Apo (red) and in the Ca^2+^-bound (blue) states. **(A)** Root-mean-square deviation (RMSD) of CALX-CBD12 backbone atoms with respect to the first frame; (**B**) Relative angle between CBD1 and CBD2; (**C**) Backbone root-mean-square fluctuation (RMSF) of CALX-CBD12 in the Apo (red) and Ca^2+^-bound states (blue) calculated over 0.4 s up to 3.4 *μ*s. The grey shaded areas correspond to the loops of the, *β*-sandwich: (I)AB-loop, (II)BC-loop, (III)CD-loop, (IV)DE-loop, (V)EF-loop, (VI)FG-loop; (**D**)Crystallographic model of CALX-CBD12 (PDB 3RB5) colored according to the relative RMSF between Apo and Ca^2+^-bound states (RMSF_*Apo*_-RMSF_*bound*_/RMSF_*Apo*_), where red and blue correspond to regions displaying decreased or increased fluctuations in the Ca^2+^-bound states relative to the Apo state, respectively.

Comparison of the backbone RMSF along the two trajectories indicated that Ca ^2+^-binding to CALX-CBD1 restricted the backbone motions in the Ca^2+^-binding region, corresponding to the AB, CD and EF loops of CBD1, and the linker between the two domains (Figs. 3C and 3D). Particularly larger fluctuations were observed at the inter-strand loops located opposite to the inter-domain interface, AB, CD, and EF in CBD2, and BC, DE, and FG in CBD1. The only exception is the FG-loop in CBD2, which displayed large fluctuations despite being located near the inter-domain interface. Fluctuations of the EF and FG loops in CBD2 were not affected by occupation of the Ca^2+^ binding sites in CBD1 (Figs. 3C and 3D). However, the CBD2 CD loop, despite of its location opposite to the Ca^2+^-binding sites, displayed increased flexibility in the Ca^2+^-bound state than in the Apo state. We attempted to investigate whether this observation could be in agreement with previously published NMR data for CALX-CBD12 (51), however NMR signals in this region were broad and relaxation rates could not be measured in the Ca^2+^-bound state. These observations are consistent with the presence of dynamics at the microsecond to millisecond time scale in the CBD2 CD-loop. Indeed, this loop does not show electron density in the crystallographic model of CALX-CBD12 (Fig. 3D).

Residues E455 and D552, which coordinate Ca^2+^ at sites Ca1, Ca2 and Ca3 in CBD1 (42), remained ∼4.3 Å apart during most of the trajectory in the Ca^2+^-bound state (Fig. 4). Furthermore, the orientations of D552 and D550 side chains, which coordinate Ca^2+^ at sites Ca1 and Ca2, also remained stable (**Fig. S4 in the Supplementary Material**). These observations indicate that the Ca^2+^ coordination geometry was preserved during the simulation. In contrast, in the Apo state simulation, E455 and D552 became further apart and sampled two major conformational states, indicating that the Ca^2+^-coordination geometry was disrupted and suggesting greater flexibility at the Ca^2+^-binding sites (Figs. 4 and **S4**). This conclusion is consistent with the observation that R584 was more mobile in the Apo state than in the Ca^2+^-bound state simulation (**Fig. S5 in the Supporting Material**). R584 could play a pivotal role in the stabilization of the extended inter-domain arrangement seen in the starting X-ray structure, since it participates in a hydrogen-bonding network with N615 in CBD2, and D552 that coordinates Ca^2+^ at sites Ca1 and Ca2 in CBD1 (Fig. 4). Furthermore, the formation of a salt bridge between R584 and D517 in the CBD1 EF-loop, and a hydrogen bond between D521 in CBD1 EF-loop and R673 in *α*-helix H2 (Figs. 4 and **S5**), may help to stabilize the extended arrangement between CALX-CBD1 and CALX-CBD2. These two interactions break in more compact inter-domain arrangements, in which E521 makes a new hydrogen bond with R681 in the C-terminal end of *α*-helix H2 (Figs. 4 and **S5**). Indeed, these analyzes are consistent with the observation that CALX-CBD12 deviated from the initial structure and sampled compact inter-domain arrangements during most of the trajectory in the Ca^2+^-bound state (Fig. 3). The crystallographic structure of CALX-CBD12 shows hydrophobic contacts (< 4 Å) between F519 in the EF-loop with L677 and S678 in *α*-helix H2 (Fig. 4) (42). When these contacts were examined during the course of the MD simulations it was found that the distance between F519 and S678 displayed broader bimodal distribution in the Apo state relative to the Ca^2+^-bound state (Fig. 4), consistent with the fact that CALX-CBD12 sampled more frequently a wider range of different inter-domain arrangements in the former state.

**Figure 4:**
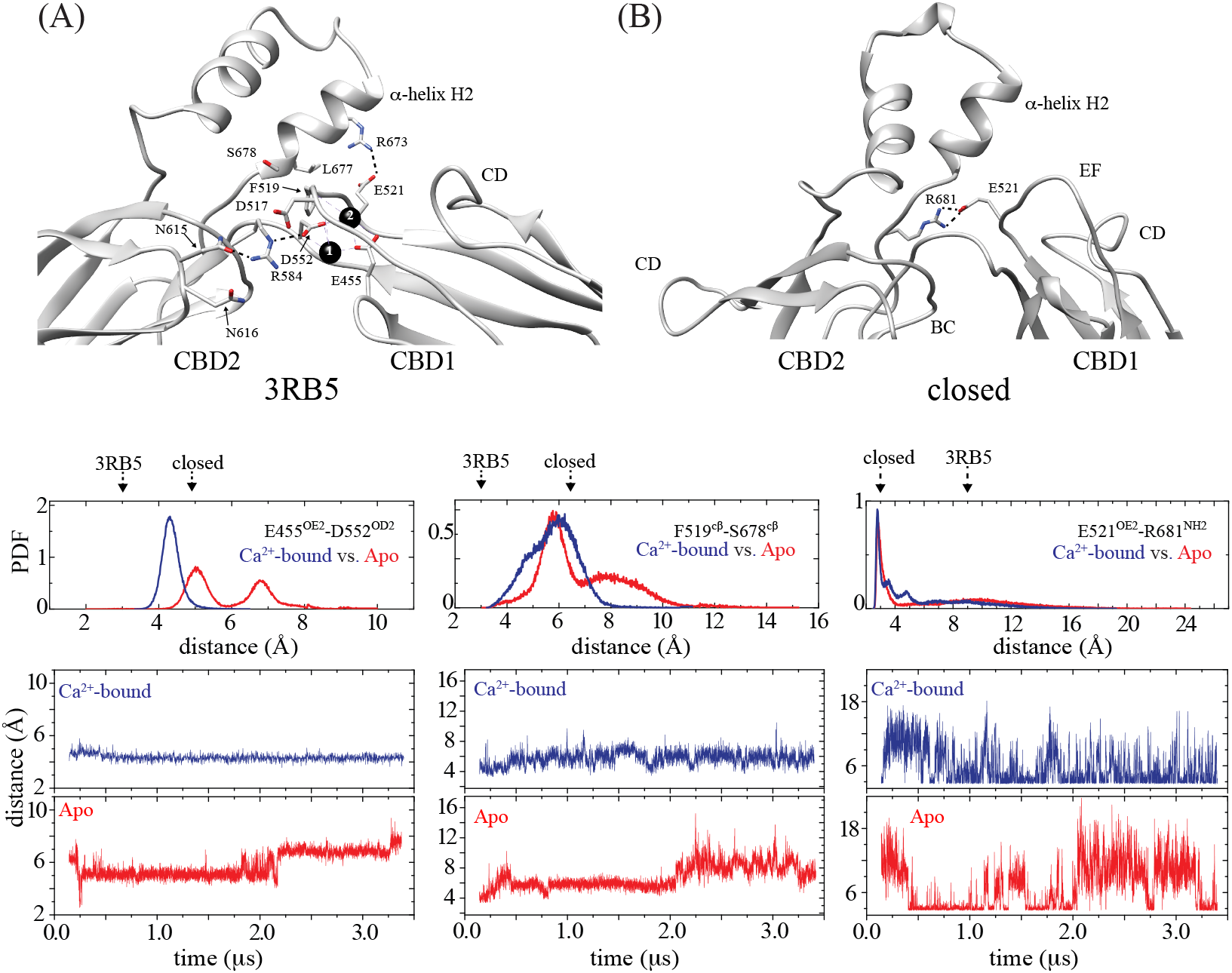
CALX-CBD12 inter-domain contacts during the MD trajectories of the Apo (red) and the Ca^2+^-bound (blue) states. **(Top)** A)Crystal structure (PDB 3RB5) of CALX-CBD12 showing the hydrogen bonding network formed between N615, R584 and D552, and the hydrogen bond formed between E521 and R673. Hydrogen bonds are indicated by dashed lines. D552 and D455 coordinate Ca^2+^ ions at Ca1 and Ca2. A possible salt bridge between D517 and R584, and a hydrophobic contact between S678 in the FG-loop of CBD2 and F519 in the EF-loop of CBD1 are also shown. B) MD snapshot of the Apo state simulation showing the formation of hydrogen bonds between R681 and E521. **(Bottom)** Pair distribution functions (PDF) and fluctuation over time of the distances between E455-D552 (**left**), F519-S678 (**middle**), and E521-R681 (**right**). Distances were computed in the interval between 0.13-3.4 *μ*s. As a reference, distances observed in the crystal structure, and in the Apo state representative snapshot are indicated by 3RB5 and “closed”, respectively.

### Using the MD trajectory as initial pool for SAXS-EOM analysis

We investigated whether refinement of an MD-derived pool against SAXS data could improve the agreement between experimental and calculated SAXS intensities. To this end, an external pool corresponding to 15 000 MD snapshots was selected and refined against SAXS data using EOM. The refined ensemble (MD-EOM) yielded a better agreement with experimental SAXS profiles than the rigid body ensemble (RANCH-EOM) as indicated by the *χ*^2^, which decreased from 6.6 to 5.2 and from 10.3 to 1.2 for the Ca^2+^-bound and the Apo states, respectively (**Figs. S2 and S6 in the Supporting Material**). The Apo state MD-EOM ensemble consisted of four *R*_*g*_ distributions, with representative models displaying *R*_*g*_ values of 23.7, 24.9, 24.9, and 26.8 Å and inter-domain angles of 75, 88, 89, and 127°. With the exception of the latter, all of them are more compact than the X-ray structure that displays *R*_*g*_ = 28 Å and inter-domain angle of 131° (**Table S1 in the Supporting Material**). It is, therefore, clear that a trend towards compact conformations is observed for the Apo state (**Fig. S6**). In the case of the Ca^2+^-bound state, two narrower *R*_*g*_ and _*max*_ distributions were obtained (**Fig. S6**). The two representative conformers displayed similar inter-domain arrangements as indicated by inter-domain angles of 126 and 129 °, consistent with a relatively rigid two-domain protein in the extended conformation (**Fig. S6**). It is noteworthy that the Apo state SAXS refined ensemble contained a population of the wide opened conformation characterized by *R*_*g*_ ∼ 26.7 Å, which is however enriched in the Ca^2+^-bound ensemble. Comparison of the initial MD pool with the MD-EOM ensemble shows that CALX-CBD12 became trapped in a local minimum far from the native structure along most of the MD trajectory in the Ca^2+^-bound state (**Fig. S6**).

Residual dipolar couplings (RDCs) measured on protein samples weakly aligned in the magnetic field report on the average orientation of the inter-nuclear bond vector with respect to an alignment tensor. Hence, RDCs are rich sources of information on structure and dynamics of proteins in solution. It was previously reported that ^1^H-^15^N RDCs obtained for CALX-CBD12 in the Apo state did not agree with the crystallographic structure as indicated by R = 0.57 obtained for RDCs measured in the presence of Pf1 (42, 51). However, better agreement was obtained with the coordinates of the separate domains as indicated by R = 0.26 and 0.34 for CALX-CBD1 and CALX-CBD2, respectively (**Fig. S7 in the Supporting Material**). Fitting the experimental RDCs to the population averaged coordinates of the four Apo state MD-EOM representative CALX-CBD12 conformers (Eqs. 2 and 3) improved the R-factor only slightly from 0.57 to 0.44 (**Fig. S6 and Fig. S8 in the Supporting Material**), suggesting that MD sampling of CALX-CBD12 conformational space was not sufficient, or that the SAXS data did not contain information on the local geometry of the HN bond vectors. This analysis yielded conformer populations that were highly correlated with each other, indicating that the correct fraction of each model was unreliable. In contrast, RDCs measured for the Ca^2+^-bound state were in reasonable agreement with the crystal structure (R = 0.35) (51), which is consistent with the rigidity of the Ca^2+^-bound state (**Fig S7**).

### Using RDCs to select native-like conformations sampled during MD

During the MD simulations CALX-CBD12 eventually visited conformations that are consistent with the ^1^H-^15^N RDCs. R-factor distributions of all MD conformers were calculated as described in the Methods section (Fig. 5). The Apo state R-factor distribution was centered at approximately R = 0.6 (Fig. 5, red), indicating that conformational sampling was insufficient to improve the agreement with the experimental RDCs data set. However, a few snapshots exhibited R factors in the range of 0.3-0.4, indicating consistency with the experimental data. These snapshots possibly represent conformers that are substantially populated in the native Apo state ensemble. The lowest R-factor MD snapshot corresponded to a compact inter-domain arrangement (Fig. 5, red), and reproduced well the experimental SAXS curve with *χ*^2^ = 1.6 using CRYSOL (**Fig. S9 in the Supporting Material**). Similarly, the R-factor distribution of the Ca^2+^-bound MD ensemble was centered at R = 0.4 (Fig. 5, blue), larger than that obtained with the crystallographic structure (R = 0.35), and consistent with the fact that CALX-CBD12 deviated from the Ca^2+^-bound native conformation during most of the MD trajectory, visiting more compact inter-domain arrangements (**Figure S6**). As observed for the Apo state, a few MD snapshots displayed improved agreement with the RDCs than the crystal structure as indicated by R-factors as low as 0.28. The snapshot displaying the lowest R-factor showed an overall agreement with the inter-domain arrangement observed in the crystal structure (Fig. 5B, blue), suggesting that the improved agreement with the experimental RDCs could result from local reorientations of the HN bond vectors rather than changes in the inter-domain orientation. Indeed, the local geometry overall quality improved during the simulation as indicated by an increase from 78 to 88 % of residues in the most favored regions of the Ramachandran plot in the MD snapshot relative to the X-ray structure. Similar observations were made for RDC data sets measured using either PEG (Apo state) or compressed polyacrylamide gels (Ca^2+^-bound state) as alignment medium and are shown in **Fig. S9**.

**Figure 5:**
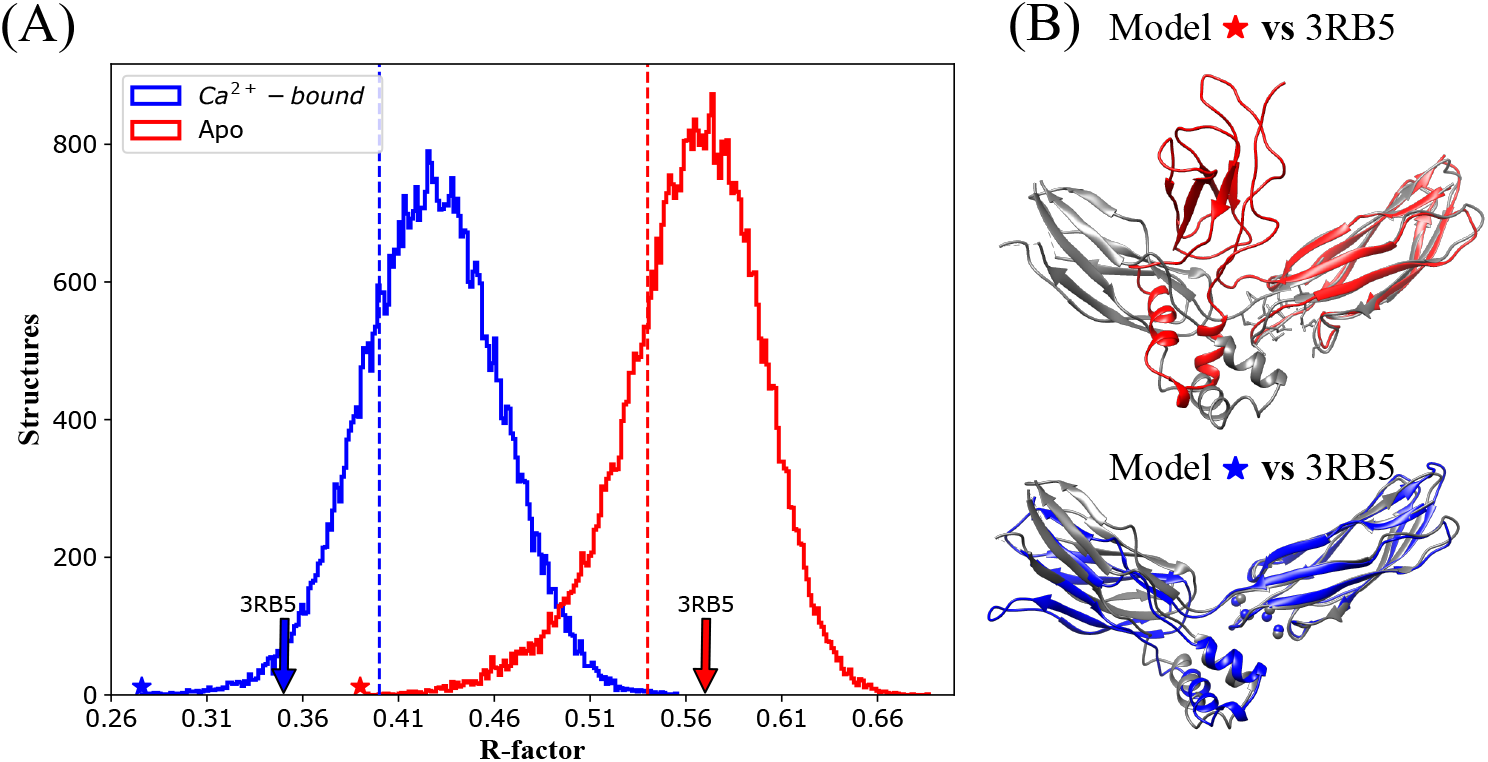
(A) RDC R-factor distributions for CALX-CBD12 MD ensembles in the Apo (red) and in the Ca^2+^-bound states (blue). Experimental ^15^N-^1^H RDCs were measured using Pf1 as alignment medium (51). MD snapshots were saved every 60 *p*s along the production phase of the trajectory (from 0.4 to 3.4 *μ*s). The vertical dotted lines indicate the R-factor cutoffs accepted to select the MD sub-ensembles used as input for EOM analyses. Arrows indicate the R-factor values obtained when CALX-CBD12 crystallographic coordinates (PDB entry 3RB5) were used for fitting. The star symbols indicate the best MD snapshots. **(B)** MD snapshots displaying best agreement with the RDCs obtained for the Apo (red) and the Ca^2+^-bound states (blue) superimposed on the crystallographic model (PDB entry 3RB5) in grey.

### Combining SAXS and RDCs to refine the CALX-CBD12 MD ensemble

In order to select a minimum ensemble that agrees with both low-resolution and high-resolution structural information, we used RDCs and SAXS experimental data to refine the MD ensemble of CALX-CBD12. We first filtered the external MD pool based on the R-factor limit of 0.54, corresponding to 15 000 snapshots (Figs. 5 and **S1**). This RDC-filtered pool was then refined with SAXS data using the EOM approach. The refined MD-RDC-EOM sub-ensemble reproduced the experimental SAXS intensities with *χ*^2^ = 1.2 and displayed a tri-modal pattern (Fig. 6). Three representative conformers were selected displaying compact (population *p*_1_ = 59 %, *θ* = 79°), semi-opened (*p*_2_ = 33 %, *θ* = 85°), and wide-opened (*p*_3_ = 8 %, *θ* = 111°) inter-domain arrangements, the latter being similar to the Ca^2+^-bound state. Additional SAXS-EOM runs were carried out starting from smaller Apo state external pools assuming lower R-factors thresholds: 0.53 (10 000 models), 0.50 (5 000 models), 0.47 (1 000 models), 0.43 (100 models) and 0.42 (50 models) (**Table S1**). After each run, the EOM representative models were re-evaluated according to the RDCs R-factor. SAXS-refinement of smaller initial pools (10 000 and 5 000 models) converged to similar sub-ensembles predominantly composed by wide-opened (*R*_*g*_= 26 Å and *θ* = 111°) and compact (*R*_*g*_= 24 Å and *θ* = 79°) models with no improvement on the population averaged R-factor, ∼ 0.43 (**Table S1 in the Supporting Material**). The information on the wide-opened conformation (*R*_*g*_ = 27 Å and *θ* = 111°) was lost when initial pools with less than 1000 conformers were submitted to SAXS-refinement by EOM, without affecting the population averaged RDC R-factor. When the flexibility of CALX-CBD12 was quantified based on SAXS data, values of *R* _*f lex*_ = 69 % and *R*_*flex*_ = 1.1 were obtained for the Apo state (**Table S1**). Despite the smallest sub-ensembles have a reduced *R* _*flex ex*_ (*R* _*flex*_ decreases from the from 78 % for the largest pool to 56 % for the smallest pool) the ratios of the standard deviations of the selected distributions with respect to the pool,*R*_*σ*_, remained greater than 0.8, near the maximum expected average of 1.0 for a fully flexible system (70) (**Table S1**).

**Figure 6:**
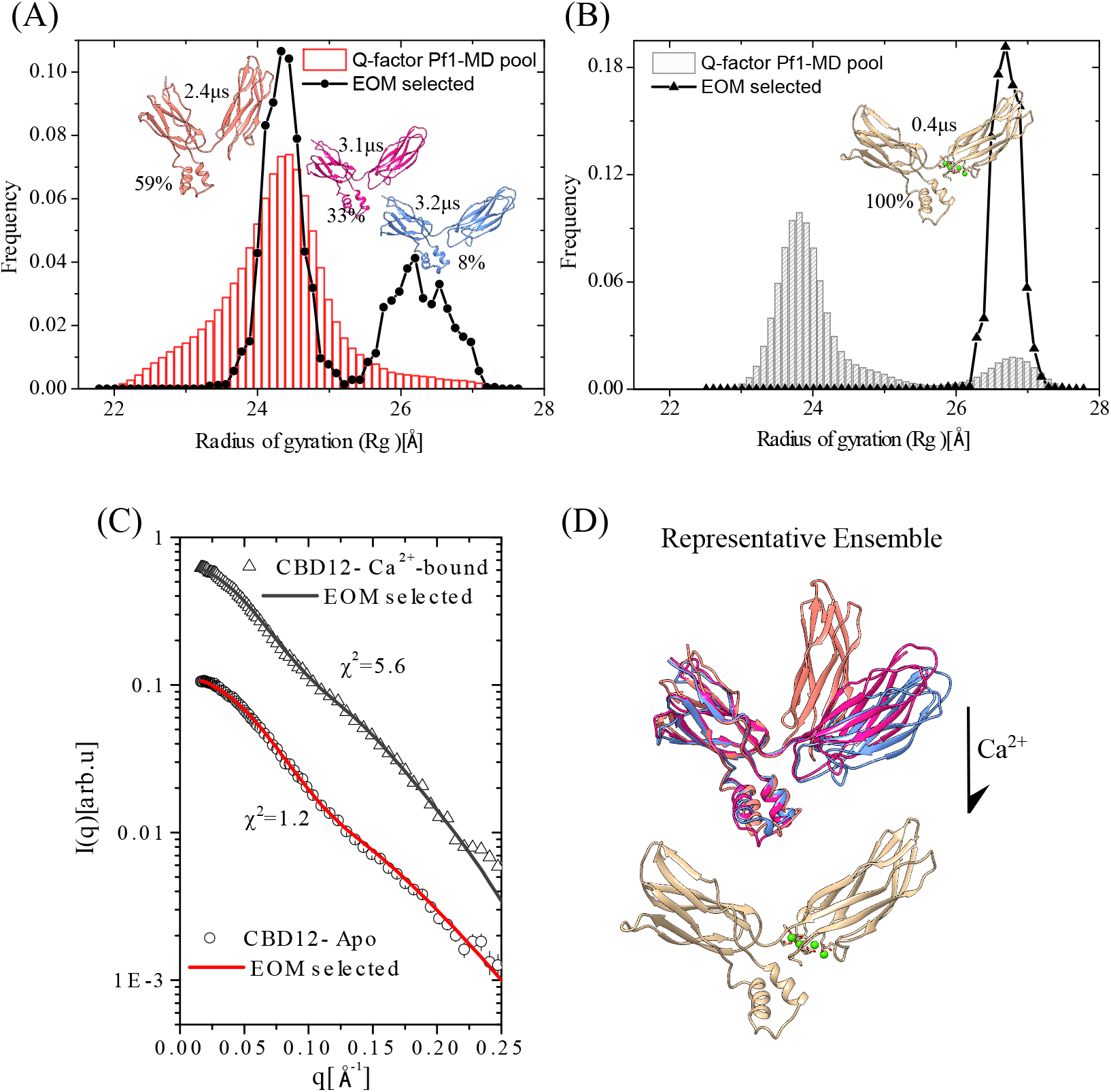
EOM selection of sub-ensembles from a pool of 15 000 MD snapshots previously filtered based on the RDCs R-factor distributions. **(A)** and **(B)** *R*_*g*_ distributions calculated for the external pool (bar graph) and the SAXS refined sub-ensemble (solid line). The representative conformers, their populations, and the MD time interval of each snapshot are indicated. **(C)** EOM fits of the experimental SAXS profiles to the selected ensembles. **(D)** Representative models of the Apo and the Ca^2+^-bound CALX-CBD12 sub-ensembles superimposed on the coordinates of the CBD2 domain.

The same MD-RDC-EOM analysis was carried out for the Ca^2+^-bound state using R-factor threshold of 0.40 corresponding to a MD ensemble of 15 000 snapshots. When the external ensemble was refined using SAXS data, the selected sub-ensemble reproduced the SAXS pattern with *χ*^2^= 5.6 and yielded a single and narrow *R*_*g*_ distribution with a representative structure displaying inter-domain arrangement (*θ* = 121°) similar to the Ca^2+^-bound state crystal structure (*θ* = 131°) (backbone RMSD of 2.3 Å), and R-factor of 0.37, nearly the same as that displayed by the crystallographic structure (R = 0.35) (Fig. 6). Flexibility analysis based on SAXS data yielded *R* _*f lex*_ = 52 % and *R*_*σ*_ = 0.12 for the Ca^2+^-bound state, consistent with a rigid system (**Table S1**). Similar results were obtained when other ^1^H-^15^N RDC data sets, measured either in PEG (Apo state) or in compressed polyacrylamide gels (Ca^2+^-bound state), were used for filtering (**Figure S10 in the Supporting Information**).

## CONCLUSION

SAXS investigations in the present work support previous NMR studies demonstrating that CALX-CBD12 behaves as a flexible two-domain protein in the Apo state, while Ca^2+^-binding shifts the equilibrium towards a relatively rigid and extended arrangement between the two CBDs. Understanding the Ca^2+^ regulation mechanism in atomic detail requires description of the various conformational states visited by CBD12 in the Apo state. Here this question was addressed using molecular dynamics simulations, SAXS and RDCs experimental data in combination with the EOM approach to select minimal conformational ensembles. It was found that CALX-CBD12 may adopt considerably closed inter-domain arrangements in the Apo state, inter-converting between three major populated conformers with wide-opened (*θ* ∼ 111 Å), semi-opened (*θ* ∼ 85 Å) and closed (*θ* 79 Å) inter-domain arrangements. In contrast, a single extended inter-domain orientation was selected in the presence of Ca ^+^. While the MD simulations sampled a range of open and closed conformations both in the Ca^2+^-bound as well as Apo state, a preference for more closed conformations was observed. Especially for the Ca^2+^-bound case this is not consistent with the experimental result of a dominating extended inter-domain orientation. One needs to consider that small inaccuracies of the force field representation by 1-2 kcal/mol are equivalent to relative population changes (e.g between closed vs open states) of a factor 10. Hence, in the absence of an extensive experimental characterization it is not possible to conclude that a set of the selected MD snapshots do represent the native conformational ensemble of CALX-CBD12 in the Apo state. However, they do highlight motions experienced by the Apo state as inter-conversions between wide-open, semi-open and closed conformations, with the equilibrium shifted towards the latter. Thus, binding of Ca^2+^ shifts this equilibrium towards the wide-open state. One may envision that in the full-length exchanger this population shift event will trigger a tension on the linker domains that eventually could be relayed to the transmembrane helices causing the inactivation of the CALX. The overall strategy of combining optimized conformation sampling from MD simulations, high resolution information from NMR-RDC data and low-resolution information from SAXS data can be a good guide for the investigation of other types of multi-domain proteins.

## AUTHOR CONTRIBUTIONS

MFSD collected and analyzed SAXS data and analyzed RDC data. PAMV prepared protein samples required for SAXS data collection. LAA provided RDCs data. MFSD and MZ carried out MD simulations and analysis of MD trajectories. MFSD, MZ, MS, RKS, and CLPO designed the research and wrote the paper.

## ACKNOWLEDGMENTS

This work was supported by grants from the São Paulo Research Foundation (FAPESP 2016/07490-1; 2019/19968-1; 2018/16092-5; 2016/24531-3) and Conselho Nacional de Desenvolvimento Científico e Tecnológico (CNPq 420490/2016-7). PMV received FAPESP PhD fellowship (2017/05614-8). MFSD and LAA received PhD fellowships from Coordenação de Aperfeiçoamento de Pessoal de Nível Superior (CAPES, #88882.160170/2014-01 and #33002010017P0). RKS and CLPO receive CNPq research fellowships (#311807/2017-8, #303001/2019-4). This work was supported by the Instituto Nacional de Ciência e Tecnologia em Fluídos Complexos, INCT-GFCx.

## SUPPLEMENTARY MATERIAL

An online supplement to this article can be found by visiting BJ Online at http://www.biophysj.org.

