## Supplemental Information for "Molecular insights on CALX-CBD12 inter-domain dynamics from MD simulations, RDCs and SAXS"

Manuscript submitted to **Biophysical** *Journal*

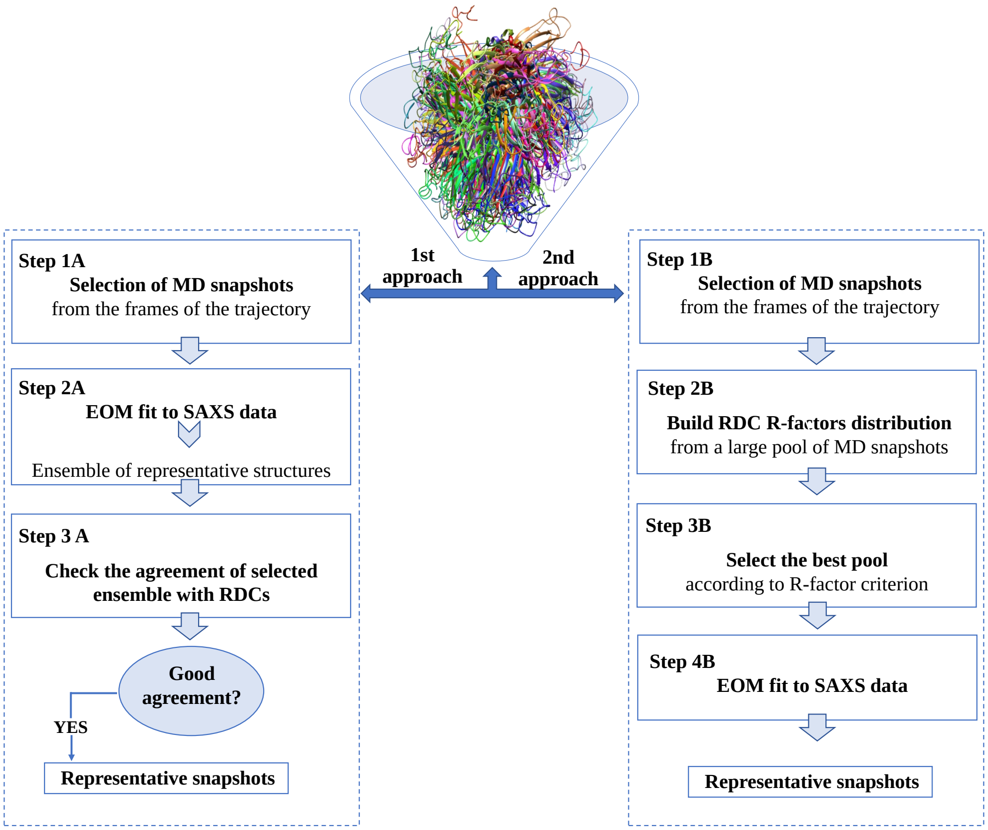

Figure S 1: **Strategy adopted to combine MD with SAXS and RDC information to build a minimal CALX-CBD12 ensemble**. A structural library was built from snapshots of the 3.4 $\mu$s MD trajectory. This initial ensemble was refined using only SAXS ($1^{st}$ approach - left) or RDC and SAXS data ($2^{nd}$ approach - right). **Left:**The initial pool derived from the MD trajectory (Step 1A) was refined against SAXS data using the ensemble optimization method (EOM). A sub-ensemble of structures fitting the SAXS data was selected (Step 2A). The agreement between the representative ensemble and the $^{15}$N-$^{1}$H RDCs data set was evaluated (Step 3A). Right: Experimental RDC data were first used to refine the initial MD pool (Step 2B), subsequently a sub-pool was selected based on the R-factor criterion and used as EOM input (Step 3B) generating a RDC-SAXS refined sub-ensemble (Step 4B).

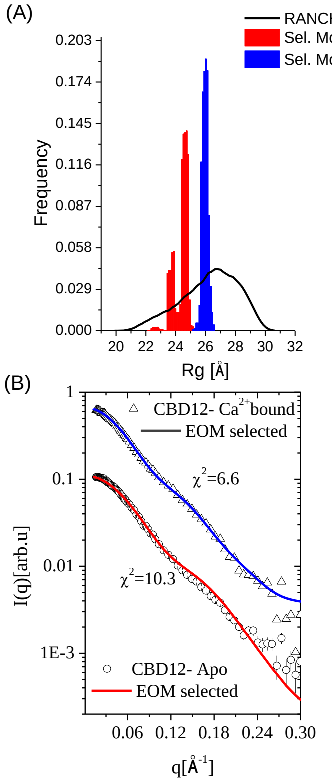

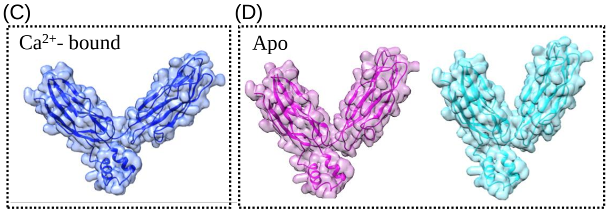

**EOM - representative structures**

| | Model | $R_g$ | $D_{max}$ | Fraction |
|---|---|---|---|---|
| $Ca^{2+}$-bound | Blue | 26 | 87.30 | 1.0 |
| Apo | Cyan | 23.76 | 78.64 | 0.7 |
| | Purple | 24.68 | 77.23 | 0.2 |

Figure S 2: **EOM-fitting of CALX-CBD12 SAXS intensity profiles to an external pool of models built by rigid-body modeling. (A)**Distributions of radius of gyration ($R_g$) computed for the external (black) and refined RANCH-EOM pools for the Apo (red) and the $Ca^{2+}$-bound states (blue). **(B)** Experimental versus fitted SAXS profiles. **(C)** and **(D)** EOM-representative models are shown in cartoon with their respective $R_g$, $D_{max}$ and population described in the table.

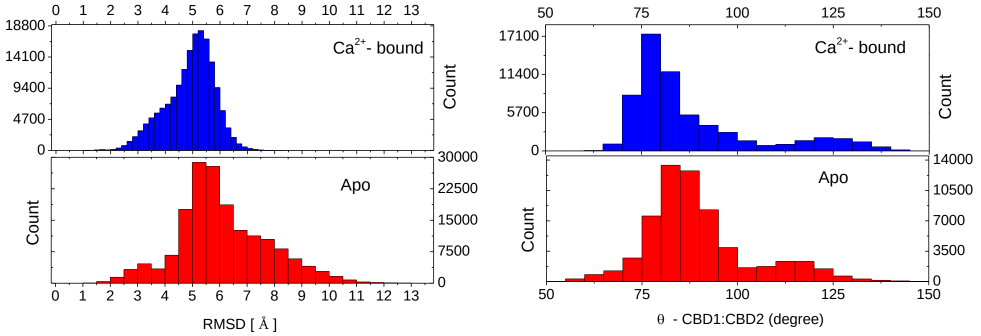

Figure S 3: **Analysis of molecular dynamics trajectories of CALX-CBD12 in the Apo (red) and in the $Ca^{2+}$-bound (blue) states. Left:** Distributions of the root-mean-square deviation (RMSD) of backbone atoms with respect to the first frame; **Right:** Distributions of the CBD12 inter-domain angle during the production phase of the trajectory.

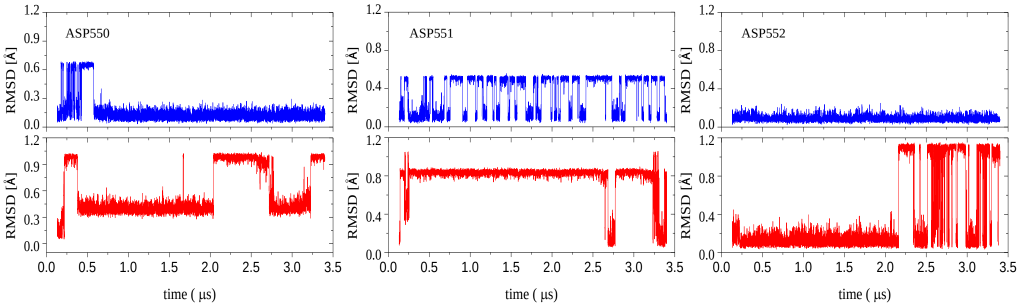

Figure S 4: **Investigation of the flexibility of D550, D551 and D552 that participate in $Ca^{2+}$ coordination at sites Ca1 and Ca2.** Sidechain RMSD with respect to the frame at 0.1$\mu$s, as a function of the simulation time.

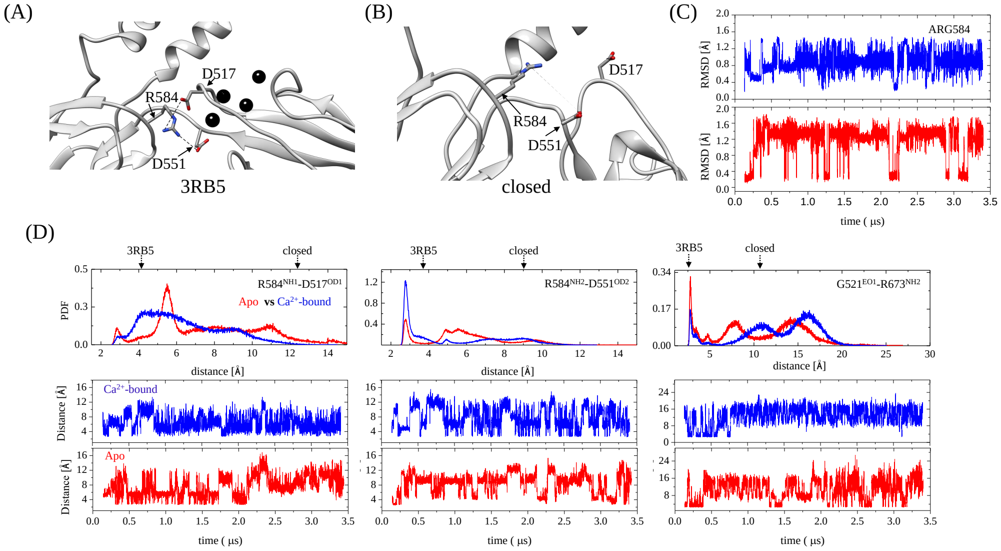

Figure S 5: **Investigation of contacts made by R584 at the interface between CBD1 and CBD2 during the MD trajectory in the Apo (red) and in the $Ca^{2+}$-bound (blue) states.** **(A)** Crystal structure of CALX-CBD12 (PDB:3RB5) showing the salt bridges formed between D517-R584 and R584-D551 at the interdomain interface. **(B)** New orientations assumed by D517, R584, and D551 in a closed conformation MD snapshot. **(C)** R584 side-chain RMSD.**(D)** Pair distribution functions (PDF) and fluctuation over time of the distances between D517- R584, D551 - D584 and G521 - R673.

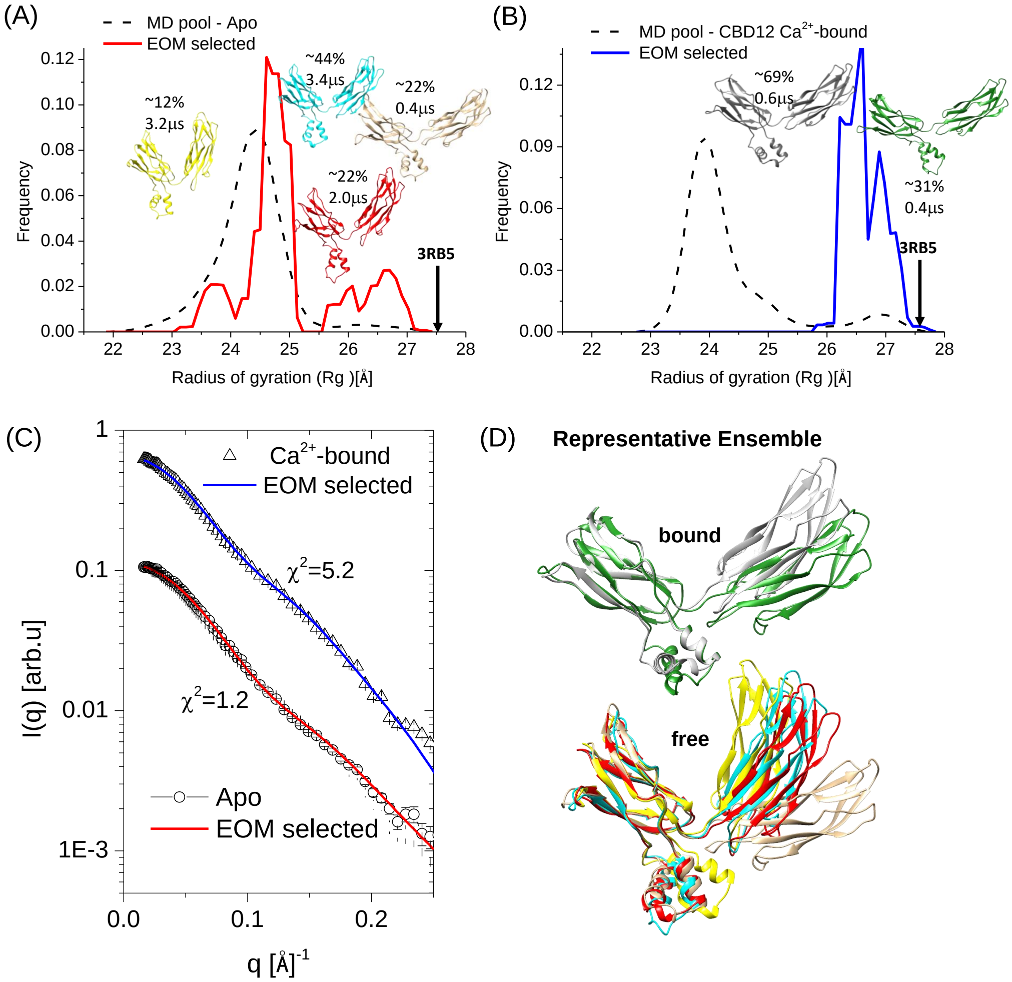

Figure S 6: **EOM-fitting of CALX-CBD12 SAXS intensity profiles to external pools derived from the MD trajectories (MD-EOM ensemble). Above**: $R_g$ distributions of the external pools of MD snapshots (dotted line) and of the refined MD-EOM ensembles (solid line) for the Apo **(A)** and $Ca^{2+}$-bound states **(B)**. Representative structures of the selected ensemble fractions are shown in cartoon with the corresponding fraction populations indicated. $R_g$ and $D_{max}$ values of the CALX-CBD12 crystallographic structure (PDB 3RB5) are indicated by arrows. **Below**: **(C)** EOM fitting of experimental SAXS profiles obtained for CALX-CBD12 in the Apo and in the $Ca^{2+}$-bound states; **(D)** Representative models are shown in cartoon superimposed on the coordinates of the CBD2 domain.

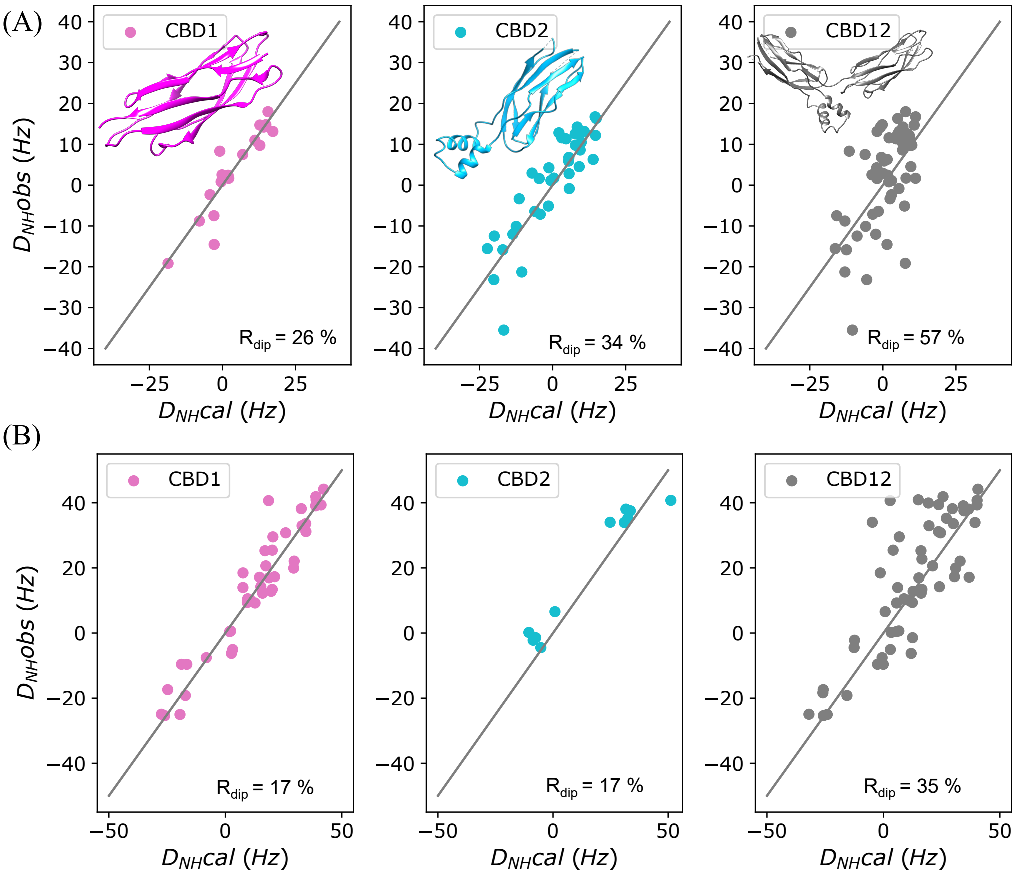

Figure S 7: Fitting of $^{1}H$-$^{15}N$ RDCs measured on CBD12 aligned with Pf1 phages in the absence (Apo, A) and in the presence of $Ca^{2+}$($Ca^{2+}$-bound, B) to the crystallographic coordinates of CALX-CBD12 or to the coordinates of the individual domains (PBD 3RB5) using Singular value decomposition (SVD).

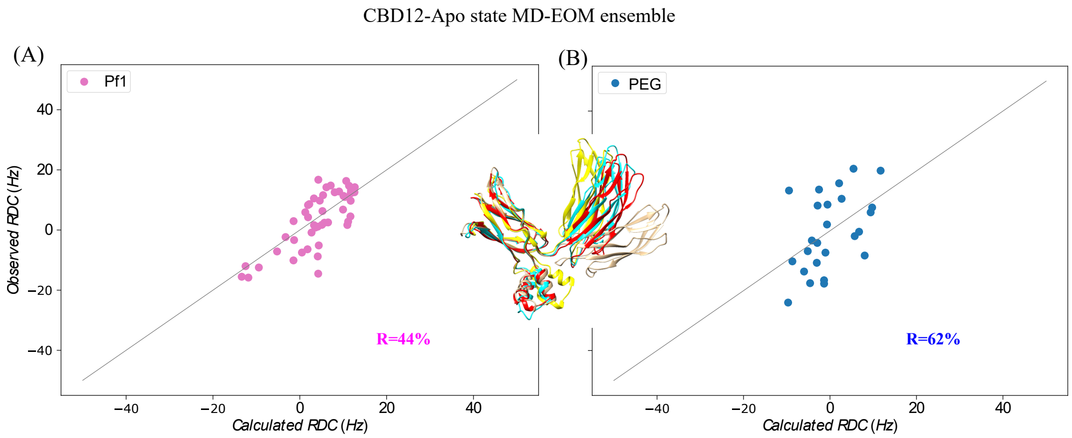

Figure S 8: **Correlation between experimental and calculated $^{1}$H-$^{15}$N RDCs**. Experimental RDCs were measured on CALX-CBD12 in the Apo state using Pf1 (left) or PEG (right) as alignment medium. The RDCs were back calculated using the average N-H bond vectors coordinates of the representative models of the MD-EOM ensemble.

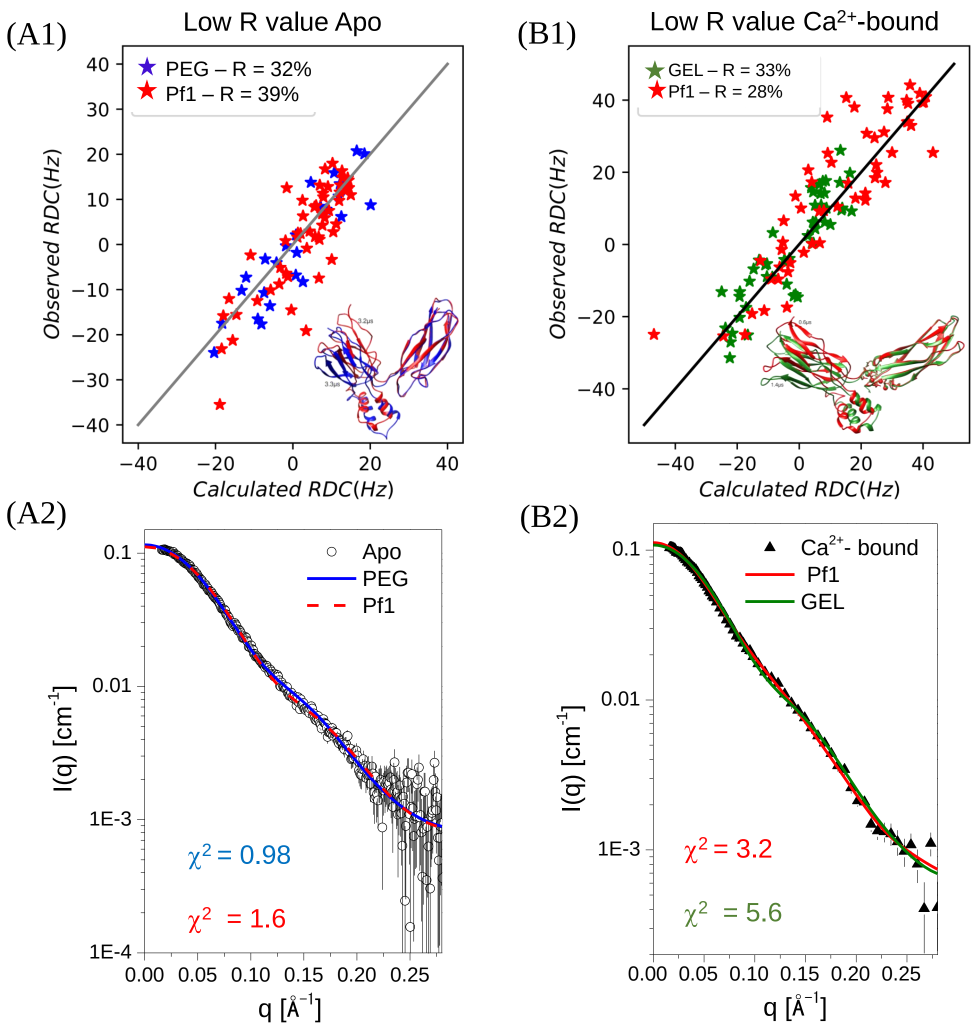

Figure S 9: **Fittings of SAXS and RDCs data to CALX-CBD12 MD snapshots displaying the lowest R-factors for experimental RDCs in the Apo state (A1 and A2) and in the $Ca^{2+}$-bound state (B1 and B2)**. Experimental RDCs were measured in PEG (blue) and Pf1 (red) for the Apo state, or in compressed polyacrylamide gel (green) and Pf1 (red) in the $Ca^{2+}$-bound state and were previously reported (1).

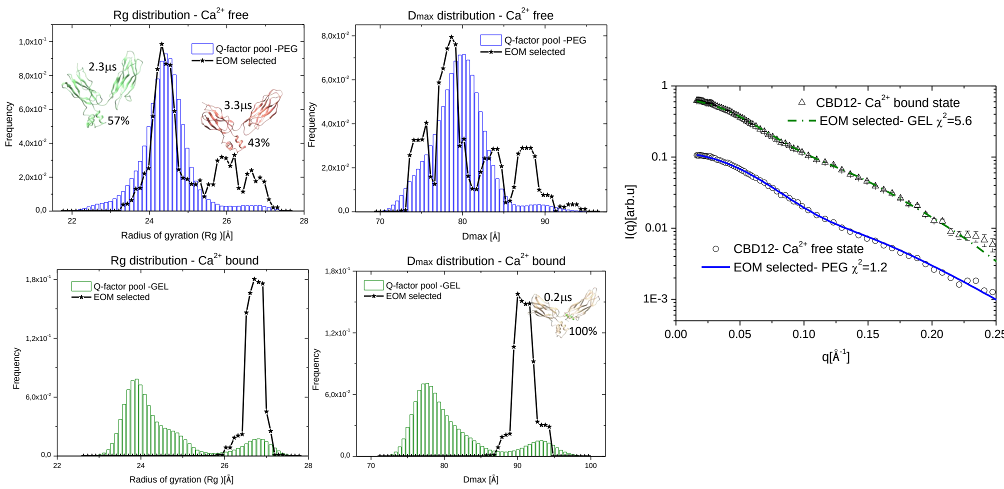

Figure S 10: **EOM selection of sub-ensembles from a pool of 15 000 MD snapshots filtered using $^{1}$H-$^{15}$N RDCs**. The RDCs were measured using either PEG (Apo state) or compressed polyacrylamide gels ($Ca^{2+}$-bound). **Left:** Panel with $R_g$ and $D_{max}$ distributions calculated for the pool (bar graph) and the optimized selected ensembles (black solid lines). The representative models selected by EOM are shown in cartoon and the corresponding simulation time is indicated. **Right:** EOM fits of the experimental SAXS profiles to the selected ensemble.

Table S 1: SAXS-based flexibility assessment of the selected MD-RDC-EOM ensembles of CALX-CBD12 in the Apo and in the $Ca^{2+}$-bound states.

| | EOM – Apo state | | | | | | | RDCs$^{b}$ |
|---|---|---|---|---|---|---|---|---|
| Pool No. of models$^{a}$ | Representative models (id/$\mu$s) | $R_g$(Å) | $\theta$(°) | Fraction | $R_g/D_{max}$ (Å) | Rflex(%) $R\sigma^{c}$ | $\chi^2$ | R(%) |
| 15 000 | apo35413 / 2.4<br>apo53068 / 3.1<br>apo53212 / 3.2 | 24.4<br>26.0<br>26.5 | 79<br>85<br>111 | 0.59<br>0.33<br>0.08 | 25.1/<br>81.0 | 69 (84)<br>1.1 | 1.2 | 44 |
| 10 000 | apo35413 / 2.4<br>apo38250 / 2.5<br>apo53212 / 3.3 | 24.4<br>24.3<br>26.0 | 79<br>81<br>111 | 0.44<br>0.11<br>0.44 | 25.1/<br>81.0 | 78 (86)<br>1.1 | 1.2 | 43 |
| 5 000 | apo35413 / 2.4<br>apo38250 / 2.5<br>apo53212 / 3.3 | 24.4<br>24.3<br>26.0 | 79<br>81<br>111 | 0.44<br>0.11<br>0.44 | 25.1/<br>81.0 | 71 (90)<br>0.97 | 1.2 | 43 |
| 1.000 | apo47341 / 3.0<br>apo53203 / 3.3 | 23.7<br>25.5 | 76<br>92 | 0.32<br>0.68 | 25.1/<br>82.0 | 62 (92)<br>0.88 | 1.2 | 41 |
| 100 | apo47341 / 3.0<br>apo53203 / 3.3 | 23.7<br>25.5 | 76<br>92 | 0.32<br>0.68 | 24.9/<br>82.8 | 56 (90)<br>0.87 | 1.4 | 38 |
| 50 | apo36574 / 2.3<br>apo53203 / 3.3 | 23.4<br>25.5 | 73<br>92 | 0.3<br>0.7 | 24.9/<br>82.9 | 56 (90)<br>0.86 | 1.4 | 37 |
| | EOM-$Ca^{2+}$-bound | | | | | | | |
| 15.000 | Ca3885 / 0.4 | 26.7 | 121 | 1.0 | 26.7 / 91.1 | 52 (81)<br>0.12 | 5.6 | 37 |
| Crystallographic structure | | | | | | | | |
| 3RB5 | 3RB5 | 28 | 131 | ——— | | | | 35 |

$^{a}$Different R-factor thresholds were adopted to select an initial EOM pool (fist column);
$^{b}$ $^{1}$H-$^{15}$N RDCs were measured for CALX-CBD12 weakly aligned with Pf1 phages;
$^{c}$The Rflex of the selected ensemble is compared to that of the initial pool in round brackets;
